# A knockout screening platform for interferon-stimulated genes in respiratory cells identifies LY6E as a dominant antiviral effector against coronaviruses

**DOI:** 10.64898/2026.09.21.751853

**Authors:** Caitlin I. Stoddard, Morgan L. Litchford, Kevin Sung, Rebecca L. Hutcheson, Joshua Marceau, Taylor Johnson, Akinobu Ota, Nell E. Baumgarten, Michael Gale, Julie Overbaugh

**Author notes:** Present address: Department of Microbiology and Immunology, and the Institute on Infectious Disease, University of Minnesota, Minneapolis, Minnesota, USA. Present address: Department of Food and Nutritional Environment, College of Human Life and Environment, Kinjo Gakuin University, Nagoya, Japan (Akinobu Ota).

## Abstract

Human coronaviruses (HCoVs) cause pathogenic outcomes ranging from mild illness to severe respiratory disease. Determining how respiratory cells coordinate an early antiviral response is necessary to understand successful control of HCoV replication. The Type I Interferon (IFN) response is a major component of innate viral immunity. IFN signaling leads to the upregulation of hundreds of interferon-stimulated genes (ISGs) that can have antiviral activity, though the individual ISGs that are responsible for restricting mildly pathogenic HCoVs in the respiratory epithelium are not clear. Here, we developed a targeted CRISPR-Cas9 knockout sgRNA library (termed the respISG library), focusing on ISGs that are upregulated across a panel of respiratory cells, to screen for innate host factors that restrict the mildly pathogenic human coronavirus, HCoV-OC43. We executed cell death-based screens in immortalized human small airway epithelial cells and identified lymphocyte antigen 6E (LY6E) as a top hit. We confirmed LY6E activity in validation studies and found it regulates entry of various SARS-CoV-2 Spike-pseudotyped lentiviruses. In addition to demonstrating a key role for LY6E in blocking a variety of HCoVs in small airway epithelial cells, this study establishes a loss-of-function CRISPR-Cas9 screening platform for identification of HCoV antiviral ISGs in respiratory cells.

**Importance:** Interferon-stimulated genes (ISGs) are upregulated during the innate immune response and contribute to early antiviral defense. High-throughput genetic screening approaches to identify functional ISGs have revolutionized our understanding of interferon-mediated control of several viruses, including coronaviruses. However, screens that focus on cells most relevant for replication of the target virus tend to be less common. Here, we develop a loss-of-function screen based on ISGs from a panel of respiratory cell models and use it to probe antiviral genes that target HCoV-OC43 in human small airway epithelial cells. We find LY6E, a well-known viral effector, to be a dominant antiviral ISG and demonstrate that its activity, in a cell type that has broad relevance to HCoV pathogenesis, also extends to SARS-CoV-2 variants.

## Introduction

Three human coronaviruses (HCoVs) with high pathogenicity have emerged and caused major global health consequences in recent decades (SARS-CoV-1, MERS-CoV and SARS-CoV-2). Less pathogenic HCoVs, including HCoV-OC43, -229E, -NL63 and -HKU1, have circulated for longer and mainly cause mild illness (1, 2), suggesting the human immune response is better equipped to confront these viruses compared to their lethal counterparts that emerged more recently. How the innate immune response, particularly the Type I Interferon (IFN) pathway, interferes with the replication of less pathogenic HCoVs is not entirely clear. During the Type I IFN response, cells sense viral nucleic acid and initiate the production of IFN, a cytokine that stimulates the upregulation of hundreds of IFN-stimulated genes (ISGs) in infected and bystander cells (reviewed in (3)).

Intact IFN signaling is necessary for innate immunity against HCoVs (4–6), but the full complement of ISGs that are responsible for IFN’s protective effect across respiratory compartments is not fully understood.

High-throughput functional genetic screens, including CRISPR-Cas9 approaches and cDNA screens, have expanded understanding of host effectors involved in HCoV infection. Prior screens have identified proviral host factors that are required for HCoV replication (7–13), while just a few have investigated antiviral host effectors. Of these screens targeting antiviral genes, genome-wide strategies have been employed (9, 12, 14) but were not designed to detect ISGs unless they are also expressed basally. Screens focused specifically on ISGs include two using overexpression approaches (15, 16), and one using a CRISPR knockout (KO) approach that targeted ISGs upregulated in cells relevant to HIV infection rather than cells that are critical to HCoV replication in infected individuals (17). Thus, while progress has been made to define host factors that control HCoV infection, targeted ISG KO screens to assess the antiviral potential of ISGs specifically in respiratory cells are needed.

To better understand the ISGs that restrict mildly pathogenic HCoVs, we sought to develop a customized functional screen with high relevance to IFN-mediated immunity in respiratory viral infection. We opted for a loss-of-function strategy to probe ISG function at endogenous levels, rather than an overexpression approach, which may skew protein concentrations away from physiological conditions. Here, we describe a respiratory ISG-focused (“respISG”) CRISPR-Cas9 sgRNA library that we applied to immortalized human small airway epithelial (HSAEC1-KT-ACE2) cells to identify ISGs that restrict the mildly pathogenic HCoV, HCoV-OC43. Our screen identified lymphocyte antigen 6E (LY6E), previously reported as both an antiviral and a proviral effector (9, 12, 14–23), as a potent restrictor of HCoV-OC43 in cells derived from the human small airway epithelium. We validate LY6E’s activity using single-gene CRISPR-Cas9 KO in HSAEC1-KT-ACE2 cells and demonstrate that it restricts various SARS-CoV-2 variants in these cells. Together, our study establishes a new screening pipeline to identify antiviral ISGs relevant to respiratory infection and highlights the importance of LY6E in restricting HCoV infection in a small airway epithelium cell model.

## Results

### Development of a targeted respiratory cell-specific ISG knockout library

To develop a customized respiratory cell-focused sgRNA library targeting ISGs for CRISPR-Cas9 KO, we used bulk RNA sequencing to identify genes upregulated upon Type I IFN treatment in three respiratory cell models (HSAEC1-KT-ACE2, Calu3 and H1299) and referenced a previously published ISG dataset from IFN-β-treated A549 cells (24). After 24-hours, a nearly identical set of genes was upregulated among all cell lines treated with IFN-α compared to IFN-β (Supplementary Table S1), including several canonical ISGs such as members of the Mx, OAS and IFIT families (Fig. 1A and B). Calu3 cells exhibited an attenuation in ISG induction overall (Fig. 1A), while HSAEC1-KT-ACE2 cells were highly responsive to Type I IFN treatment, exhibiting over 200 unique ISGs that were not upregulated in the other three cell lines (Fig. 1B). There were 102-139 genes in common across any two cell types, with fewer shared across three cell types (between 58-96 genes). Only 57 genes were ISGs across all four cell lines (Fig. 1B).

**Figure 1.**
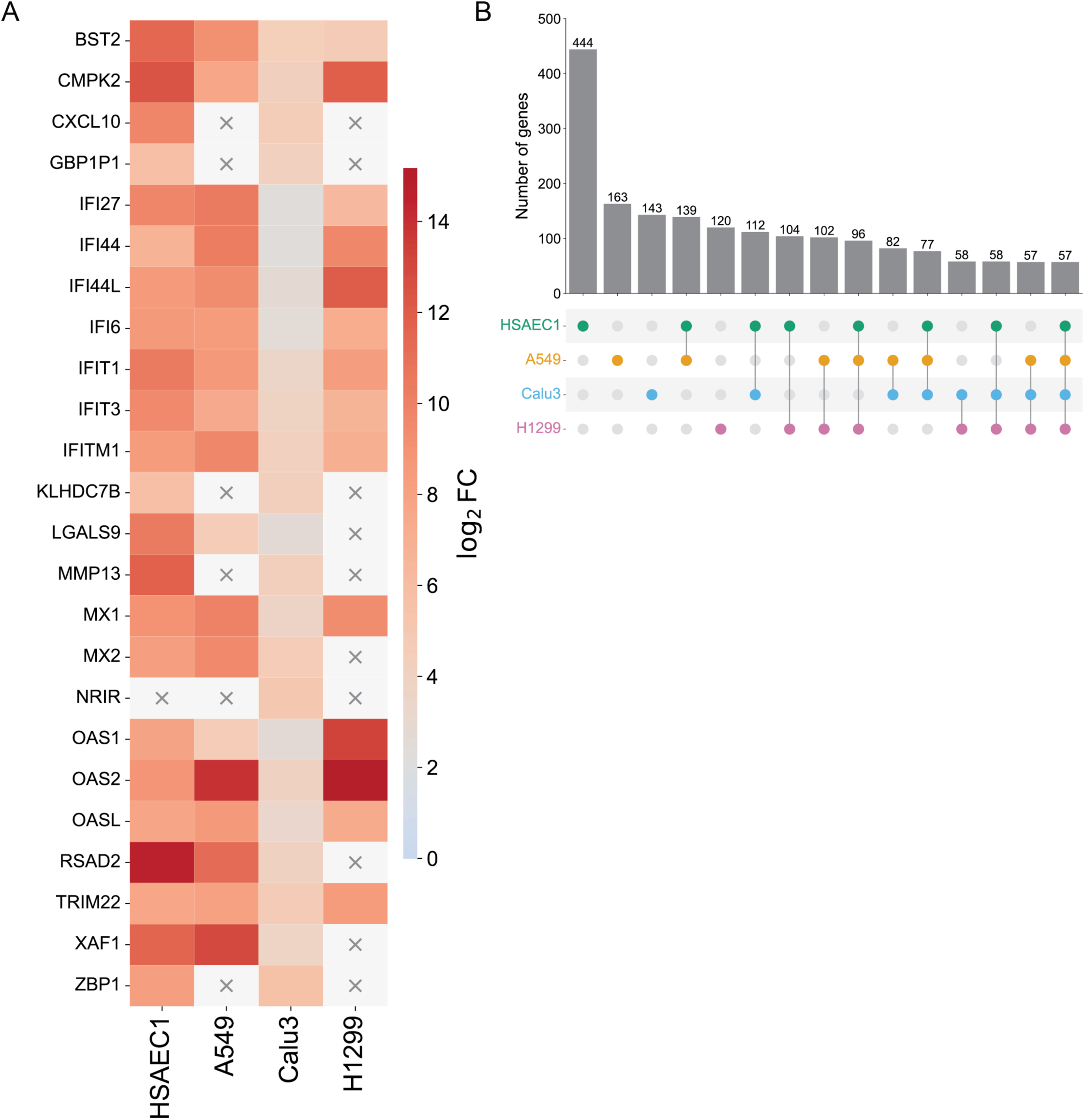
Bulk RNA-seq to identify interferon-stimulated genes in respiratory cell models. (A) Heatmap showing differential expression (DE) (log_2_FC) for the top DE genes for each cell line treated with IFN-β. Genes that were not induced in each cell line are indicated with an “X”. (B) UpSet plot depicting intersections of IFN-β-induced DE genes shared across HSAEC1-KT-ACE2, A549, Calu3, and H1299 cells.

We selected ISGs that were differentially expressed (DE) upon IFN treatment compared to untreated cells in any of the cell lines for inclusion in a sgRNA library for use with CRISPR-Cas9. We also included a subset of genes that were upregulated by SARS-CoV-2 infection in HSAEC1-KT-ACE2 or Calu3 cells (Supplementary Tables S1 and S2) since they have the potential to contribute to antiviral effects. After filtering out genes that were not protein-coding, the library included sgRNAs targeting 562 respiratory cell-specific DE genes plus 31 additional genes related to previous screens conducted with SARS-CoV-2 (23) or other viral families (24, 25) (Table 1). The final library, which we call the “respISG” library, contained 192 non-targeting control (NTC) sequences, 622 candidate genes, and 5152 total sgRNAs (Table 1).

**Table 1.**
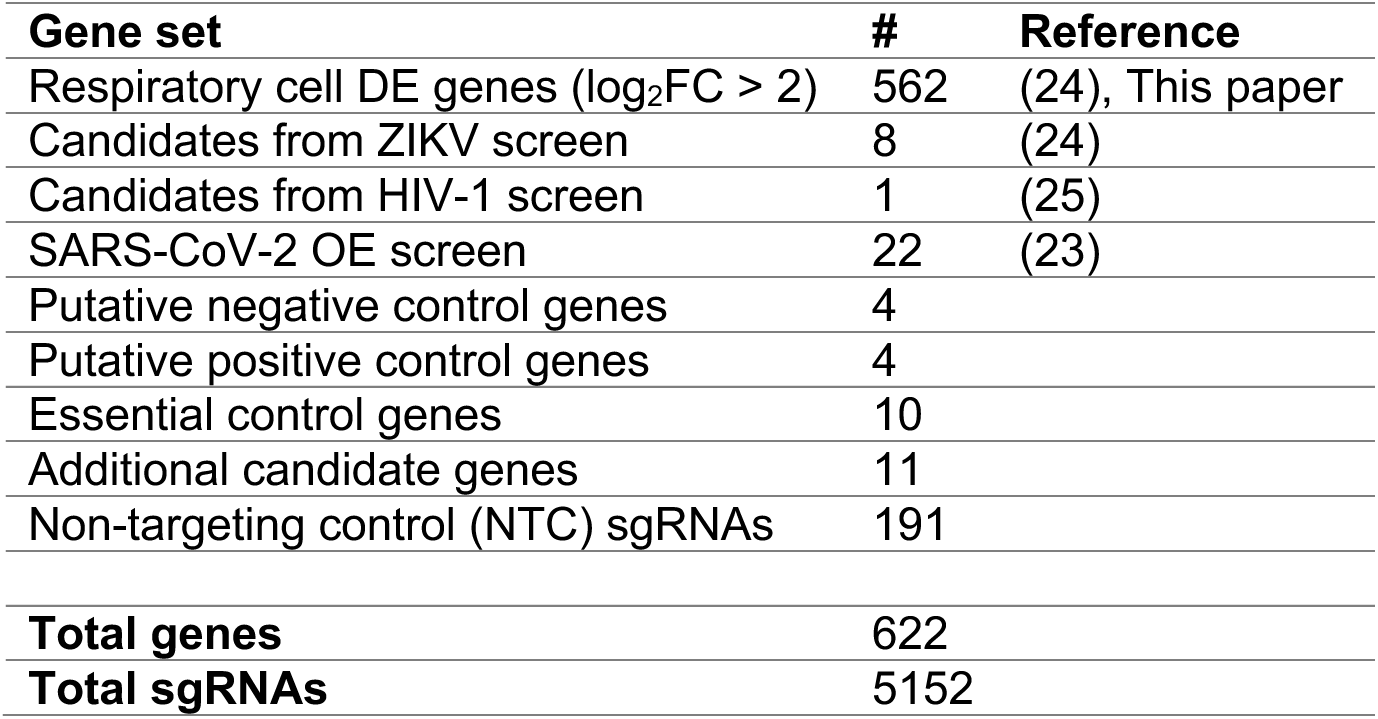
RespISG sgRNA library summary. Differentially expressed (DE) genes from this study were combined with genes identified as antiviral candidates in screens for Zika virus, HIV-1, or SARS-CoV-2 as indicated. Additional genes with no known antiviral function (putative negative control genes) or genes related to HCoV entry (putative positive control genes) were included along with a subset of essential genes and non-targeting control (NTC) sequences. The final number of genes and sgRNAs in the respISG library is indicated.

| **Gene set** | **#** | **Reference** |
| --- | --- | --- |
| Respiratory cell DE genes ($\log_2\text{FC} > 2$) | 562 | (24), This paper |
| Candidates from ZIKV screen | 8 | (24) |
| Candidates from HIV-1 screen | 1 | (25) |
| SARS-CoV-2 OE screen | 22 | (23) |
| Putative negative control genes | 4 |  |
| Putative positive control genes | 4 |  |
| Essential control genes | 10 |  |
| Additional candidate genes | 11 |  |
| Non-targeting control (NTC) sgRNAs | 191 |  |
| **Total genes** | **622** |  |
| **Total sgRNAs** | **5152** |  |

### CRISPR-Cas9 knockout screen to identify anti-HCoV-OC43 ISGs in human small airway epithelial cells

To identify ISGs with antiviral activity against HCoV-OC43, we assessed IFN-mediated protection during HCoV- OC43 infection of HSAEC1-KT-ACE2, Calu3 and MRC-5 cells (Supplementary Fig. S1A-C). IFN-β treatment most robustly protected HSAEC1-KT-ACE2 cells during HCoV-OC43 infection, with a greater than 3000-fold reduction in virus replication after 78 hours at an MOI of 0.1 or 0.01, and these cells were selected for downstream screening (Supplementary Fig. S1A). We found that HCoV-OC43 infection of HSAEC1-KT-ACE2 cells induced cell death (Supplementary Fig. S2), which allowed us to design a CRISPR-Cas9 screen using cytopathy as a functional readout akin to our recently reported screen with Zika virus (24) (Fig. 2A).

**Figure 2.**
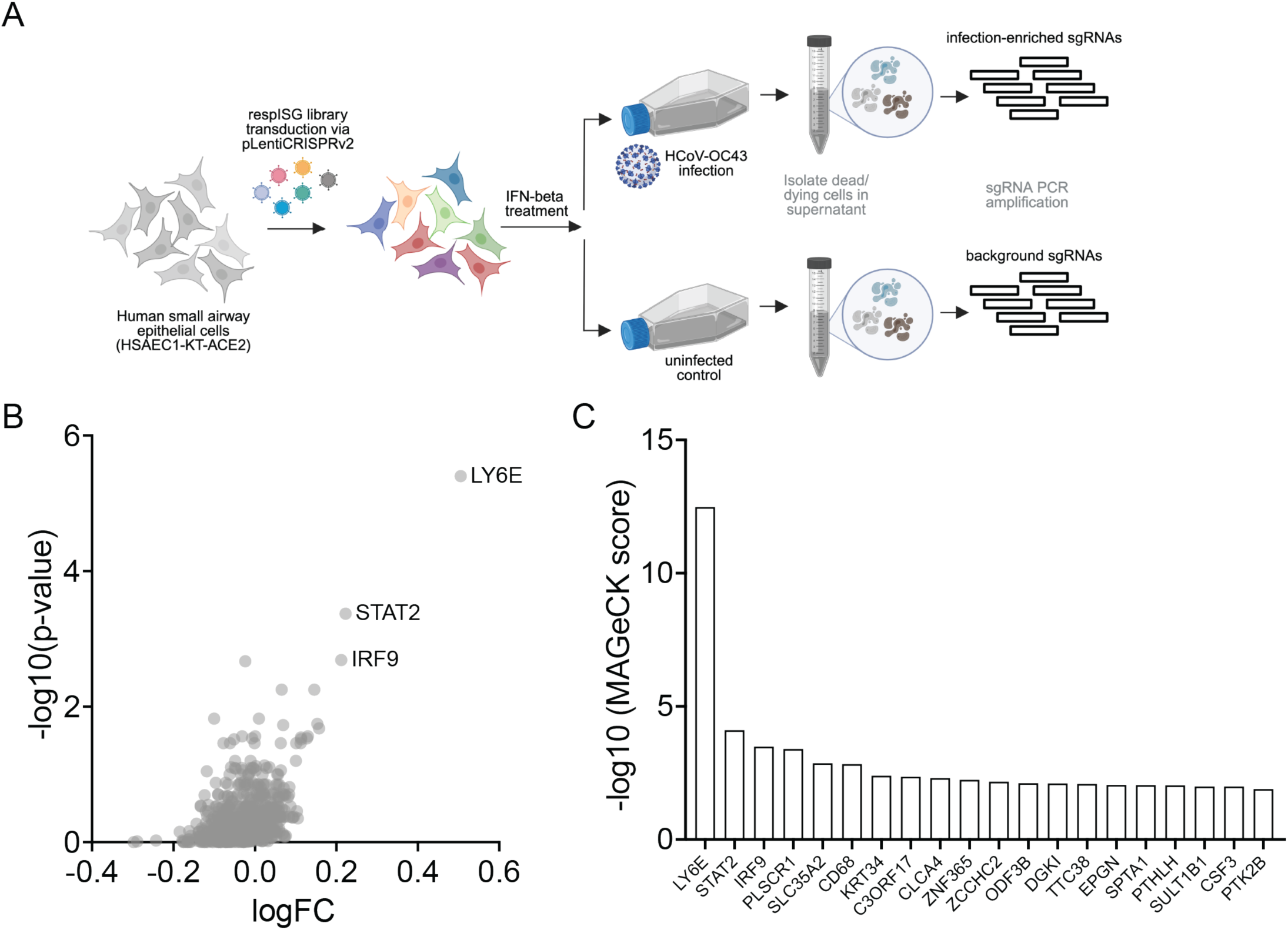
CRISPR-Cas9 respISG cell death screen for antiviral genes that restrict HCoV-OC43. (A) HSAEC1- KT-ACE2 cells were transduced with a lentiviral library containing respISG sgRNAs and Cas9. Library cells were treated with IFN-β (500 U/mL) and left uninfected or infected with HCoV-OC43 at an MOI of 0.5. Dead and dying (detached) cells were isolated from the culture supernatant after 2, 3 or 4 days. sgRNAs were amplified and sequenced from control and infected populations and sgRNAs enriched in the infected population were evaluated as candidate antiviral genes. (B) Gene enrichment in infected cells across three combined harvest days (2, 3 and 4 days post-infection) calculated using MAGeCK software with NTC normalization. (C) MAGeCK scores for genes corresponding to sgRNAs enriched in infected cells across the three harvest days. The top 20 genes are depicted.

Using this approach, the screen enriched for STAT2 and IRF9, both regulators of ISG transcription, highlighting that the platform could successfully identify genes relevant to IFN-mediated protection against HCoV-OC43 (Fig. 2B). Notably, LY6E, a gene with anti-HCoV activity first reported in human hepatoma and A549 lung adenocarcinoma cells (16, 22), was the most enriched hit in the screen, surpassing STAT2 and IRF9 in MAGeCK score and p-value (Figs. 2B and C). PLSCR1, a gene previously shown to be antiviral in hepatoma (26, 27) and A549 (27, 28) cells infected with SARS-CoV-2, was the next highest enriched gene in the screen. Given its dominance in the screen, we focused on validation of LY6E as an anti-HCoV ISG in HSAEC1-KT-ACE2 cells.

### LY6E blocks HCoV entry in human small airway epithelial cells

To determine whether LY6E was antiviral against HCoV-OC43 in HSAEC1-KT-ACE2 cells outside the context of the screen, we independently generated two HSAEC1-KT-ACE2 LY6E KO cell pools and confirmed LY6E KO via Western blot (Fig. 3A). As expected, LY6E expression was low in NTC cells in the absence of IFN treatment and was stimulated by IFN-β (Fig. 3A). We infected untreated and IFN-β-treated NTC, IRF9 KO, and LY6E KO cell pools with HCoV-OC43 and measured HCoV-OC43 Nucleocapsid levels using RT-qPCR. IFN-β treatment of NTC cells reduced HCoV-OC43 replication throughout the time course of infection. The antiviral effect was more than 15-fold after 48 hours at an MOI of 0.5 and KO of IRF9 ablated the protective effect of IFN-β treatment in HSAEC1-KT-ACE2 cells, bringing replication levels to those of untreated NTC cells (Fig. 3B). In two independent LY6E KO cell pools, HCoV-OC43 replication was appreciably higher than in NTC cells in the presence of IFN-β (Fig. 3B), suggesting LY6E is antiviral against HCoV-OC43 in these cells. Notably, LY6E KO did not fully restore replication to the levels in cells that were not treated with IFN-β. By comparing HCoV-OC43 replication levels in LY6E KO cells to NTC and IRF9 KO cells at 48 hours post-infection, we estimate that LY6E accounts for approximately 22% of the protection conferred by IFN treatment (Fig. 3C). This observation suggests that while LY6E is a dominant antiviral gene in this system, other factors contribute to IFN-mediated protection in this cell model. Complementing LY6E KO cells with exogenous LY6E (Fig. 3D) reduced HCoV- OC43 replication back to the level observed with endogenous LY6E in NTC cells (Figs. 3E and F), confirming the antiviral effect was attributable to LY6E expression.

**Figure 3.**
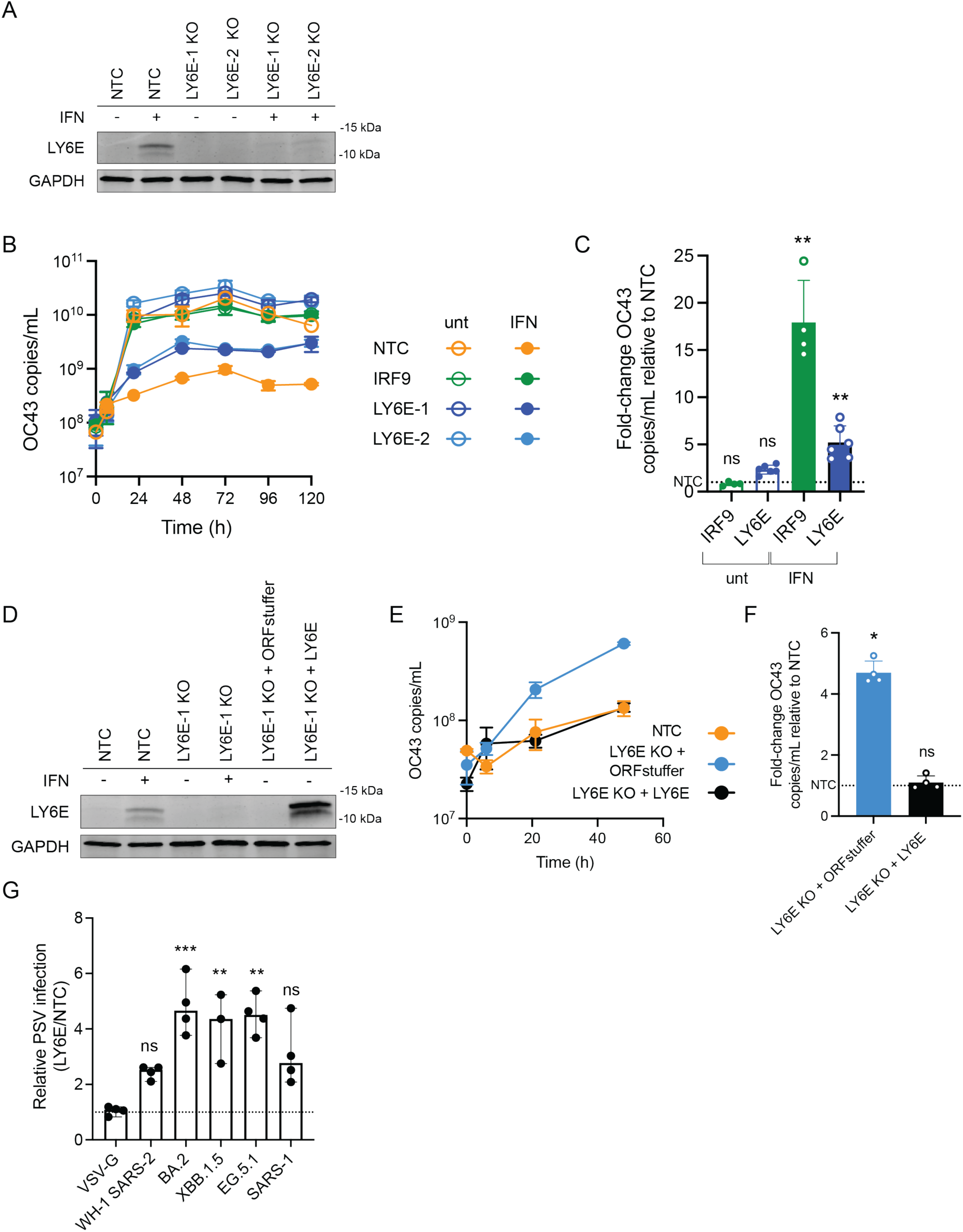
LY6E restricts HCoV-OC43 and SARS-CoV-2 variants in human small airway epithelial cells. (A) Western blot for LY6E in HSAEC1-KT-ACE2 NTC or LY6E KO cell pools in the presence or absence of 500 U/mL IFN-β. GAPDH was included as a loading control. (B) RT-qPCR for HCoV-OC43 Nucleocapsid during representative infection of NTC, IRF9 KO, or LY6E KO HSAEC1-KT-ACE2 cells in the presence or absence of 500 U/mL IFN-β. Two technical replicates are presented. (C) HCoV-OC43 Nucleocapsid copies/mL determined by RT-qPCR for IRF9 KO and LY6E KO relative to NTC HSAEC1-KT-ACE2 cells at 48 hours post-infection in the presence or absence of 500 U/mL IFN-β. Dotted line indicates NTC copies/mL. Kruskal-Wallis test was applied, ns = not significant, ** = p < 0.01. Data were from at least four independent experiments. (D) Western blot for LY6E in NTC, LY6E KO, LY6E KO with control expression vector complementation, or LY6E KO with LY6E complementation in the presence or absence of 500 U/mL IFN-β. GAPDH was included as a loading control. (E) RT-qPCR for HCoV-OC43 Nucleocapsid during representative infection of NTC HSAEC1-KT-ACE2 cells, LY6E KO cells with control expression vector (ORFstuffer), or LY6E KO cells with LY6E complementation. Cells were treated with 500 U/mL IFN-β. Two technical replicates are presented. (F) HCoV-OC43 Nucleocapsid copies/mL determined by RT-qPCR for LY6E KO cells complemented with control expression vector (ORFstuffer) or LY6E at 48 hours post-infection. Cells were treated with 500 U/mL IFN-β. Dotted line indicates NTC copies/mL. Kruskal-Wallis test was applied, ns = not significant, * = p < 0.05. Data were from two independent experiments. (G) Pseudovirus (PSV) entry in untreated IFN-β-treated LY6E KO versus NTC HSAEC1-KT-ACE2 cells. X-axis labels indicate Spike entry glycoprotein used for PSV construct. The dashed line indicates a relative PSV infection value of 1 (denoting no sensitivity to PSV entry in LY6E KO vs NTC cells). Kruskal-Wallis test was applied for each Spike construct compared to VSV-G, ns = not significant, ** = p < 0.01, *** = p < 0.001. PSV data collected from at least three independent experiments.

LY6E is reported to block entry of HCoVs in hepatoma (16, 22) and African green monkey (VeroE6) cells (16, 22). To investigate whether LY6E interferes with viral entry in HSAEC1-KT-ACE2 cells, we generated a panel of pseudotyped viruses (PSVs) bearing Spikes from ancestral Wuhan-Hu-1 SARS-CoV-2, SARS-CoV-2 Omicron variants (BA.2, XBB.1.5, and EG.5.1) and SARS-CoV-1 using a lentiviral reporter system. These PSVs undergo just a single round of infection and thus allow for the isolation of the entry step of the viral replication cycle. Loss of LY6E did not impact the entry of PSVs bearing VSV-G and therefore we used VSV-G PSVs as a control for comparison to HCoV Spike-expressing PSVs (Fig. 3G). LY6E KO cells treated with IFN-β were more susceptible to entry of PSVs bearing all Spike proteins tested, ranging from 2.3- to 4.6-fold increases in infection in this single round assay, although the differences did not reach statistical significance for WH-1 SARS-CoV-2 and SARS-CoV-1 (Fig. 3G). Together, these results suggest LY6E is an IFN-dependent regulator of Spike-mediated HCoV entry, and its activity in small airway epithelial cells spans diverse HCoVs, including mildly and more severely pathogenic subgroups.

## Discussion

The identification of antiviral genes that control HCoV infection has advanced our understanding of host- HCoV interactions, particularly since the emergence of SARS-CoV-2. In addition to targeted studies of individual candidate genes, there is added value in high-throughput genetic screens as a means of systematic antiviral gene discovery. One such approach has been to identify candidate antiviral genes using genome-wide loss-of-function strategies (9, 12, 14). This approach allows for study of gene activity at endogenous levels; however, the lack of IFN treatment limits our ability to examine the effects of ISG KO during this key antiviral response. Conversely, screens designed for the identification of antiviral ISGs have largely employed overexpression strategies (15, 16, 23), or repurposed screens for other viruses (17), rather than focusing on the specific impact of respiratory ISGs at endogenous levels. The screen described here was designed to bridge these powerful strategies by creating a specific respiratory cell-focused ISG sgRNA library to conduct a KO screen in respiratory cells derived from the lower airway.

LY6E, a member of the LY6/uPAR family and the predominant hit in our screen, has been implicated historically in promoting infection with influenza A (19), flaviviruses (18) and HIV-1 (20). More recently, LY6E has emerged in several screens for HCoV antiviral host effectors in different cell and HCoV contexts (9, 12, 14–17).

LY6E’s antiviral role has been validated across several cell types including non-respiratory cells and cancerous respiratory cell lines, though not previously in the small airway epithelium. The small airway epithelium, part of the lower respiratory tract, has been implicated in HCoV infection associated with severe outcomes (reviewed in (29)). Our results reinforce LY6E’s important role in blocking HCoV infection in respiratory cells, which play an essential role in HCoV pathogenesis.

Infection with HCoVs is associated with pathogenic outcomes ranging from mild to severe. Our screen demonstrated that LY6E is antiviral against replicating HCoV-OC43, consistent with a recent HCoV-OC43 screen conducted in A549 cells, a cancerous respiratory cell line where LY6E is basally expressed (14). We were interested in whether endogenous LY6E could restrict more lethal HCoVs like SARS-CoV-2, including Omicron variants that emerged later and exhibited unique infection and transmission characteristics (30–33). Deletion of LY6E in IFN-treated cells enhanced the entry of PSVs decorated with Spikes from SARS-CoV-2 variants, suggesting LY6E antiviral activity in HSAEC1-KT-ACE2 cells extends to more lethal HCoVs. Overall, this study establishes a new respiratory cell-focused loss-of-function ISG sgRNA library for CRISPR-Cas9 screens in the context of respiratory viral infection and emphasizes the importance of LY6E in blocking infection with diverse HCoVs in a range of tissue contexts, including cells from the small airway epithelium.

## Acknowledgments

We are grateful to Vrasha Chohan, Jamie Guenthoer, and Felicitas Ruiz for assistance with pseudovirus production. Spike expression plasmids for SARS-CoV-1, XBB.1.5 and EG.5.1 were a gift from Marceline Cote. Spike expression plasmids for WH-1 and BA.2, lentivirus accessory plasmids, and 293T-ACE2 cells were a gift from Jesse Bloom. pLentiCRISPRv2 (Addgene #52961) was a gift from Feng Zhang. psPAX2 (Addgene #12260) and pMD2.G (Addgene #12259) were gifts from Didier Trono. We thank Hannah Itell for advice on sgRNA library construction and Alexandra Willcox for advice on ISG complementation. We are grateful to Frederick Matsen and members of the Overbaugh and Emerman labs at Fred Hutch for helpful discussion and advice. Thank you to the Fred Hutch Genomics core for assistance with Illumina sequencing and Pritha Chanana for assistance with bioinformatics. This work was supported by NIAID K99/R00 grant AI171000 awarded to C.I.S., NIAID grants AI183793 and AI179722 to M.G., Jr., and Endowed Chair Funding to J.O.

## Materials and Methods

### Cells and viruses

Immortalized human small airway epithelial cells (HSAEC1-KT, ATCC) were transduced with human ACE2 using a lentivirus system followed by cell sorting to generate HSAEC1-KT-ACE2 and were cultured in Airway Epithelial Cell Growth Medium (Promocell C-21060) with 1X Penicillin-Streptomycin (Gibco 15140-122). H1299 human non-small lung cancer cells (ATCC CRL-5803) were maintained in RPMI (Gibco 1640) supplemented with 10% fetal bovine serum (FBS), 2 mM L-glutamine (Gibco 25030081), and 1X antibiotic-antimycotic (anti-anti, Gibco 15240062). A549 lung adenocarcinoma cells (A. Berger; ATCC CCL-185) were cultured in RPMI supplemented with 10% FBS, 2 mM L-glutamine, and 1X anti-anti. Calu3 lung adenocarcinoma cells (ATCC HTB-55) were maintained in EMEM (ATCC 30-2003) supplemented with 10% FBS and 1X anti-anti. MRC-5 human lung fibroblast cells were cultured in EMEM with 10% FBS, 2 mM L-glutamine, and 1X anti-anti. HCT-8 (ATCC CCL- 244) colorectal adenocarcinoma cells were cultured in RPMI supplemented with 10% horse serum (Gibco 16050130). 293T (ATCC CRL-3216) and 293T-ACE2 (J. Bloom) cells were maintained in DMEM (Gibco 11965092) with 10% FBS, 2 mM L-glutamine, and 1X anti-anti. All cell lines were authenticated with STR profiling prior to experimentation.

HCoV-OC43 (Betacoronavirus 1, VR-1558) was provided by BEI. HCoV-OC43 was propagated in HCT-8 cells and viral titers were determined by TCID_50_ in MRC-5 cells. SARS-CoV-2 isolate USA-WA1/2020 was derived from BEI (NR-52281) and propagated in Vero cells (USAMRIID). SARS-CoV-2 titers were determined by plaque assay in Vero cells.

### Bulk RNA sequencing and data analysis

Bulk RNA sequencing to identify ISGs in respiratory cell lines was conducted in two separate experiments with overlapping conditions. For H1299, cells were plated in triplicate in the presence or absence of 1000 U/mL IFN- α or IFN-β. After 24 hours, cells were harvested and total RNA was extracted using the RNeasy Plus Mini Kit (Qiagen 74106). RNA quality and concentration was evaluated using RNA TapeStation (Agilent). Illumina RNA-sequencing was performed with 50 bp paired-end reads. RNA-seq analysis was conducted as described previously for A549 cells (24). Briefly, sequencing reads were mapped to the GRCh38 human reference genome using GENCODE annotations (34) via Spliced Transcripts Alignment to a Reference (STAR) 2.7.9a software (35). Gene counts were analyzed with edgeR (36) and genes were required to have at least 10 counts in some samples and 15 counts total across all samples (using filterByExpr function) to be defined as an ISG. The glmQLFTest function was used to determine if variances for the gene between IFN-treated and untreated conditions were significantly different. ISGs were required to pass the edgeR filter with p < 0.05 and have counts per million (CPM) > 1 in at least half the samples across both treated and untreated conditions. Log_2_ fold-change (log_2_FC) was calculated using the average CPM in the IFN-treated condition over the untreated condition. Analyses were conducted in Python v3.8.10 using the Numpy v1.23.5 and Pandas v1.5.2 packages.

For Calu-3 and HSAEC1-KT-ACE2, cells were treated with 1000 U/mL IFN-β or infected with SARS-CoV- 2 at an MOI of 5 for 24 hours. Total RNA was extracted from the cells using the Quick-RNA kit (Zymo R1057T). RNA-seq libraries were constructed using the KAPA mRNA HyperPrep kit (Roche KK8580). Libraries were quality controlled and quantified using RNA TapeStation (Agilent) and Illumina sequenced with 100 bp paired-end reads. Bioinformatic analyses were completed using the R statistical programming language (v4.3.1). Genes were required to have ten or more raw counts averaged across all samples to be included. Counts were normalized via Trimmed Mean of M-values (TMM) with edgeR and log2CPM calculated with the voom function in the *limma* R library (37). For IFN-β treatment or SARS-CoV-2 infection relative to untreated/uninfected cells, differential expression analysis was performed using the *lmFit* function in *limma*. For both datasets, a gene was considered differentially expressed if log2FC > 2.

### respISG CRISPR-Cas9 sgRNA library

sgRNA sequences were designed for each protein-coding ISG or control gene included in the library using CHOPCHOP(38) or GUIDES (39). For genes with fewer than eight sgRNAs, additional sgRNA sequences were included from the Broad Institute CRISPick database (40, 41) to achieve a total of eight sgRNAs/gene. A total of 191 non-targeting control (NTC) sgRNA sequences were included as negative controls. After removal of duplicates, the respISG library contained 5152 unique sgRNA sequences covering 622 genes. sgRNA sequences were synthesized (Twist Biosciences) and amplified across eight separate PCR reactions with Herculase II Fusion DNA Polymerase (Agilent 600675) using the following primers: Array F TAACTTGAAAGTATTTCGATTTCTTGGCTTTATATATCTTGTGGAAAGGACGAAACACCG; Array R ACTTTTTCAAGTTGATAACGGACTAGCCTTATTTTAACTTGCTATTTCTAGCTCTAAAAC. PCR reactions were purified (Qiagen 28106), gel extracted (Qiagen 28704), and cloned via Gibson assembly (NEB E2611L) into BsmBI-digested pLentiCRISPRv2 (Addgene #52961). The Gibson reaction was desalted by dialysis across a 0.025 µm filter (Millipore Sigma VSWP02500) and subsequently electroporated into Endura ElectroCompetent Cells (Lucigen 60242-2) using a Gene Pulser System (Bio-Rad) with the following settings: 10 µF, 600 Ohms, 1800 Volts. Electroporation reactions were plated on 245 mm square LB-agar dishes with 50 μg/mL Carbenicillin and harvested the following day by scraping the bacterial lawn and rinsing with LB medium. The final respISG pLentiCRISPRv2 plasmid library was isolated from bacteria using the Plasmid Plus Mega Kit (Qiagen 12981), amplified as described below for screen samples, and submitted for Illumina sequencing to confirm representation of all expected sgRNAs. Lentivirus was prepared by transfection of the library into 293T cells. 4E6 HSAEC1-KT-ACE2 cells were seeded across two T-150 flasks and transfected with FuGene6 (Promega E2692) plus 10 μg respISG pLentiCRISPRv2 library, 10 μg psPAX2 (HIV-based packaging vector, Addgene #12260), and 10 μg pMD2.G (VSV-G envelope vector, Addgene 12259) per flask. Virus-like particles (VLPs) were harvested 48 hours later, concentrated using 100 kDa Amicon filters (Millipore UFC9100) (3750 x g, 15 minutes, 4°C) and used to transduce the library into HSAEC1-KT-ACE2 cells.

### respISG library transduction in HSAEC1-KT-ACE2 cells

To determine the appropriate amount of VLPs to transduce to achieve roughly 1-2 pLentiCRISPRv2 vector copies per cell, the VLP stock prepared above was titrated in HSAEC1-KT-ACE2 cells and vector copies were determined using Droplet Digital PCR (ddPCR) (Bio-Rad, see below). To maintain at least 500-fold coverage of the sgRNA library, we aimed to transduce a minimum of 11E6 HSAEC1-KT-ACE2 cells. HSAEC1-KT-ACE2 cells were seeded in six 24-well dishes at a density of 5E4 cells/well. 48 hours later, a VLP master mix was prepared (13.3 μL VLPs/1E6 cells, 9 μg/mL DEAE Dextran) in complete Airway Epithelial Cell Growth Medium and applied at 1 mL per well. Cells were spinoculated at 1200 x g for 90 minutes at room temperature, washed with 1X PBS, replenished with fresh medium and incubated overnight at 37°C. Cells were allowed to reach confluency and then serially expanded before collecting a sample for gDNA isolation and freezing at -80°C in 10E6 cell aliquots. gDNA was isolated from 0.25E6 cells (Qiagen 51104) and digested with BglI (NEB R0143L) prior to ddPCR. To determine final pLentiCRISPRv2 vector copies per cell, 3 μL digested gDNA was combined with 17 μL ddPCR

Master Mix containing 10 μM primers and probe (ddPCR-F1 CGGATCGGCACTGCG; ddPCR-U6-R ATGGGAAATAGGCCCTCG; cPPT-probe 6-FAM/ZEN-AGACATAATAGCAACAGACATACAAAC-IBFQ), 1 μL RPP30 ddPCR Copy Number Variation HEX Assay (Bio-Rad 100-31243), and 10 μL SuperMix with no dUTP (Bio-Rad 1863024). After droplet generation and PCR amplification, and data acquisition was performed on an QX200 droplet reader (Bio-Rad) using the FAM/HEX with ABS protocol. Final sgRNA library vector copies per cell for respISG HSAEC1-KT-ACE2 KO cells was approximately 0.9.

### respISG CRISPR-Cas9 knockout screen

HSAEC1-KT-ACE2 respISG KO library cells were plated in the presence of 500 U/mL human IFN-β (PBL Assay Science 11410) at a density of 2E6 cells in T-175 tissue culture flasks. Sufficient flasks were prepared to account for three separate cell harvest days (2, 3, and 4 days post-infection), including both uninfected and HCoV-OC43 infected conditions. Two flasks were plated per condition, and a second screen replicate with identical conditions was plated on a separate day. Cells were incubated for 48 hours to allow for transcriptional activation induced by IFN-β treatment and were subsequently infected with HCoV-OC43 at an MOI of 0.5. At days 2, 3 and 4 post-infection, dead/dying cells were harvested from each flask by centrifuging the supernatant and collecting the pellet. Cells were washed twice in 1X PBS and stored at –80°C until further processing. gDNA was extracted from cell pellets using the QIAamp Blood Mini Kit (Qiagen 51104). gDNA samples were quantified via Nanodrop and split across the maximum number of 2 μg Round 1 PCR reactions. sgRNA-containing sequences were amplified using Herculase II Fusion DNA Polymerase (Agilent 600675) with the following primers: FWD GAGGGCCTATTTCCCATGATTCCTTCA, REV AACTTCTCGGGGACTGTGG. Round 1 PCR products were purified using the QIAquick PCR Purification Kit (Qiagen 28104) and split across four Round 2 PCRs using the following primers: FWD AATGATACGGCGACCACCGAGATCTACACTCTTTCCCTACACGACGCTCTTCCGATCTXXXXXXTCTTGTG GAAAGGACGAAACACCG, where XXXXXX is a unique indexing barcode, REV CAAGCAGAAGACGGCATACGAGATGTGACTGGAGTTCAGACGTGTGCTCTTCCGATCTTGCCACTTTTTCAAGTTGATAACGGACT. Pooled Round 2 PCRs were purified via AMPure XP Beads (Beckman Coulter A63881) and quantified via Qubit. Equimolar barcoded samples (40 ng each) were pooled and agarose gel-extracted using a QIAquick Gel Extraction Kit (Qiagen 28704) before a final Qubit quantification to determine library concentration. Multiplexed samples were submitted for single-end 50 bp NextSeq P1 Illumina sequencing (Fred Hutch Genomics Shared Resource).

### CRISPR Screen Data Analysis

Pooled samples were demultiplexed and assigned to sample IDs based on barcodes. Adapters were trimmed from sgRNAs and sequences were aligned to the respISG sgRNA library using Bowtie (42). A set of “synthetic” NTCs was generated from respISG library NTC sequences by iteratively binning eight NTCs per synthetic NTC to reach an equivalent number of genes. sgRNA enrichment in dead and dying cells collected from the culture supernatant was compared between IFN-treated, HCoV-OC43 infected and IFN-treated, uninfected populations using the MAGeCKFlute package (43). Data were analyzed from individual harvest days (2, 3, or 4 days post-infection), or all harvest days combined over two independent screen replicates. The final analysis presented in Fig. 2 includes all harvest days in duplicate.

### Single-gene knockouts in HSAEC1-KT-ACE2 cells

Single-gene KO cell pools were generated via lentiviral transduction of lentiCRISPRv2. sgRNAs (LY6E: GTGACTGTGTCTGCTAGTGC; IRF9: ACAATTCCACAGGCCAGCCA) were annealed and ligated into BsmBI digested pLentiCRISPRv2. Two independent KO cell pools were generated for LY6E. VLPs were produced by transfection of 293T cells (T-75 flask at 70-80% confluency) with FuGene6 (Promega E2692) plus 5 μg of sgRNA-containing pLentiCRISPRv2, 5 μg psPAX2, and 5 μg pMD2.G. After 48 hours, VLPs were harvested from the supernatant by removing cellular debris via centrifugation (1200 x g, 5 minutes, 4°C) and concentration using a 100 kDa Amicon filter (Millipore UFC9100) (3750 x g, 15 minutes, 4°C). For transduction, 2E5 HSAEC1-KT-ACE2 cells/well were seeded in 6-well dishes for 24 hours before application of 50-200 μL concentrated VLPs and spinoculation at 1200 x g for 90 minutes at room temperature. Transduced cells were passaged at least three times prior to collection of 3E5 cells for gDNA isolation with the QIAamp Blood Mini Kit (Qiagen 51104). Editing was confirmed via Sanger sequencing and Inference of CRISPR Edits (ICE, Synthego/EditCO) analysis. Because cells were not subjected to selection, editing was re-tested periodically and found to be stable for at least 60 days in culture.

### LY6E complementation of LY6E knockout cells

Lentiviral LY6E and control “ORF stuffer” expression vectors were designed and synthesized with VectorBuilder. VLPs were prepared and harvested as described above using 2.5 μg LY6E or ORF stuffer vectors, 5 μg psPAX2, and 2.5 μg pMD2.G per T-75 flask seeded with 2E6 293T cells. VLPs were serially diluted and HSAEC1-KT- ACE2 cells transduced in the presence of 10 μg/mL DEAE-Dextran. Once confluent, LY6E KO HSAEC1-KT- ACE2 cells were selected with 10 μg/mL Hygromycin B (InvivoGen ant-hg-1) for seven days. Following RNA extraction (RNeasy Plus Mini Kit, Qiagen 74106) and reverse transcription (Superscript III, Invitrogen 18080- 044), exogenous LY6E mRNA expression levels were confirmed using RT-qPCR with the following primer sets: LY6E Assay Set 3 FWD GATCTTCTTGCCAGTGC, REV GGCAGTACAGATTGCTCTT; LY6E Assay Set 4 FWD GTGCTTCTCCTGCTTGA, REV TAGTTGTCCTGGTCGGA. GAPDH was used as a reference gene using the following primers: FWD GAAGGTCGGAGTCAACGGATTT, REV GAATTTGCCATGGGTGGAAT.

### HCoV-OC43 infection of HSAEC1-KT-ACE2 cells and Nucleocapsid RT-qPCR

HSAEC1-KT-ACE2 cells with LY6E modifications were seeded in 24-well dishes at a density of 5E4 cells/well. After 24 hours, medium +/- 500 U/mL IFN-β (PBL Assay Science 11410) was used to replenish cells. An additional 48 hours after IFN-β treatment, cells were infected with HCoV-OC43 at an MOI of 0.5 in serum-free Airway Epithelial Cell Growth Medium. Three hours post-infection, cells were washed three times in 1X PBS before complete medium replacement +/- IFN-β. Supernatant time course samples were collected and stored at -80°C prior to heat-inactivation at 56°C for one hour. Inactivated samples were then stored at -20°C. Thawed samples were diluted 1:5 in UltraPure water and tested via RT-qPCR (iTaq Universal Probes One-Step Kit, Bio- Rad 1725140) to determine HCoV-OC43 Nucleocapsid copies/mL using an adapted primer/probe set (44). A plasmid containing the HCoV-OC43 Nucleocapsid sequence (GenBank AY585228.1) was used to create a standard curve.

### Annexin-V staining

6.5E5 HSAEC1-KT-ACE2 cells/well were plated in a 24-well dish in the presence or absence of 1000 U/mL IFN-β (PBL Assay Science 11410). 24 hours later, cells were infected with HCoV-OC43 at an MOI of 0.01 or mock-infected. Three hours after infection, medium (+/- IFN-β) was replaced. Cells were harvested and pelleted at 48, 72, 96, 120 and 144 hours after detachment with Accutase (Gibco A11105). Staining of dead cells was carried out using an Annexin-V-FITC Apoptosis Kit (BioVision K101-25) according to manufacturer’s instructions. Briefly, cell pellets were resuspended in 100 μL Binding Buffer containing 1 μL Annexin V-FITC and incubated at room temperature for five minutes in the dark. Cells were pelleted and resuspended in 100 μL fresh Binding Buffer and fixed with 100 μL 2% paraformaldehyde prepared in Binding Buffer. Annexin V-FITC staining was evaluated using flow cytometry with a BD FACSCanto II instrument. Data were analyzed using FlowJo v10 software.

### Western blotting

HSAEC1-KT-ACE2 cells with LY6E modifications were trypsinized, pelleted, washed once in PBS, and stored at -80°C until lysis in ice-cold RIPA buffer (Cell Signaling Technologies 9806) with protease/phosphatase inhibitor (Pierce A32955). Lysates were subjected to three freeze/thaw cycles consisting of a 5 sec vortex followed by freezing in a dry ice/EtOH bath. Lysates were cleared by centrifugation at max speed for 15 min at 4°C. Protein concentrations were quantified using DC Protein Assay II (Bio-Rad 5000112) against a standard curve of bovine serum albumin (BSA). Samples were diluted to equivalent concentrations and heated to 70°C for 10 minutes in Laemmli sample buffer (Bio-Rad 1610747) containing 2.5% β-mercaptoethanol (Bio-Rad 1610710XTU). 10 μg of protein extracts were resolved on 4-20% Mini-PROTEAN TGX Protein Gels (BioRad 4561094) and transferred to 0.2 μm nitrocellulose membranes (Invitrogen IB23002) using the iBlot 2 Transfer dry transfer system (Invitrogen IB21001) set to program P0 (20–25 Volts, 7 minutes). The blots were blocked in 3% BSA for 30 minutes at room temperature and probed overnight at 4°C with anti-LY6E antibody (Invitrogen PA5-143999, 1:1000). For GAPDH staining, membranes were probed with anti-GAPDH antibody (Cell Signaling Technologies, 2118S, 1:10,000). Membranes were probed with secondary antibody for 1 hour at room temperature using IRDye 800CW-linked anti-rabbit IgG (LICORbio 926-32211, 1:20,000) and visualized fluorescently on a LICOR Odyssey CLx instrument. When necessary, membranes were stripped (Thermo Scientific 46430) and re-probed with a different antibody. Membranes were washed in TBS with 0.1% Tween-20 between each step.

### Production of Spike-pseudotyped lentiviruses and PSV entry assay

Spike PSVs were prepared as previously described (45–48). Briefly, 5E5 293T cells/well were seeded in 6-well dishes. 16-23 hours later, cells were transfected with Luciferase_IRES_ZsGreen backbone, Gag/Pol, Rev, and Tat lentiviral accessory plasmids (45) and the corresponding HCoV Spike expression plasmid using FuGene6 (Promega E2692). After 25 hours, medium was replaced. 50-60 hours post-transfection, lentiviral supernatants were filtered (Steriflip, Millipore SCGP00525), concentrated (Amicon, Millipore UFC9100), and stored at -80°C. PSV stocks were titered by infecting 293T-ACE2 cells with serially diluted samples. For the PSV entry assay, black-walled, clear-bottom 96-well plates (Greiner Bio-One 655090) were seeded with 1.25E4 NTC or LY6E KO HSAEC1-KT-ACE2 cells, or 293T-ACE2 cells as a positive control in the presence or absence of 500 U/mL IFN- β (PBL Assay Science 11410) at a volume of 50 μL. PSVs produced with HCoV Spikes or VSV-G control entry protein were used to infect cells 24 hours later at a final titer of 1.5E7 RLU/mL. Uninfected, “cells only” wells were used as a negative control. 48-55 hours post-infection, 100 μL medium was removed from wells and 30 μL room temperature BrightGlo was added (Promega E2620). Relative luciferase units (RLU) were measured with a LUMIstar Omega plate reader (BMG Labtech).

---

**Supplementary Table S1.** Overlap of differentially expressed genes across IFN-α/IFN-β treatment and SARS-CoV-2 infection conditions.

| Treatment/infection condition | # DE genes ($\log_2FC > 2$) |
| --- | --- |
| IFN-β | 504 |
| IFN-α or IFN-β | 505 |
| IFN-β or SARS-CoV-2 | 739 |
| IFN-$\alpha/\beta$ or SARS-CoV-2 | 740 |

Supplementary Table S2. Log2FC across all IFN-treated or SARS-CoV-2-infected cell lines. See attached file.

**Supplementary Figure S1.**
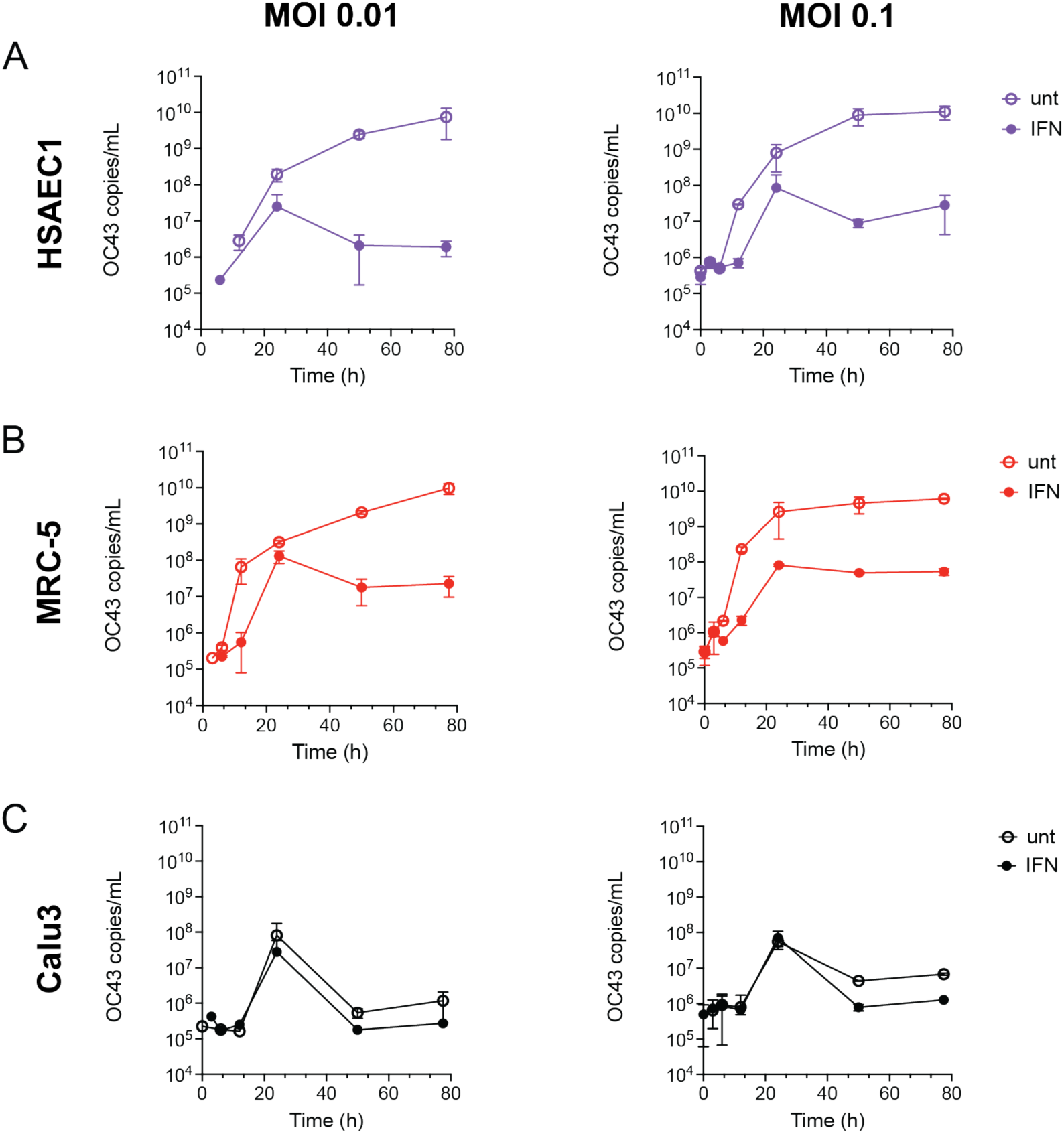
Interferon-mediated protection during infection of respiratory cell lines with HCoV- OC43 at two MOIs. (A) HSAEC1-KT-ACE2, (B) MRC-5, and (C) Calu3 cells were infected with HCoV-OC43 at an MOI of 0.01 (left panel) or 0.1 (right panel) in the presence or absence of 500 U/mL IFN-β and HCoV-OC43 Nucleocapsid copies were determined by RT-qPCR.

**Supplementary Figure S2.**
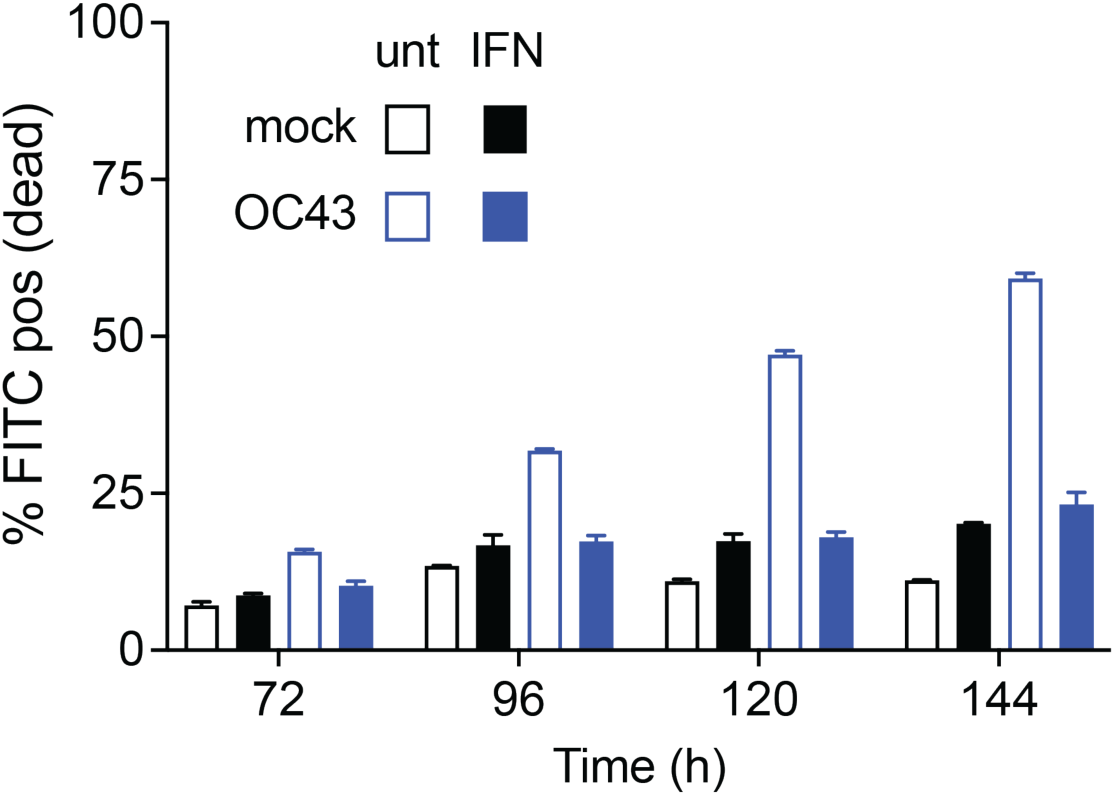
HSAEC1-KT-ACE2 cell death during HCoV-OC43 infection. Untreated or IFN-β- treated HSAEC1-KT-ACE2 cells were infected or left uninfected, stained with Annexin-V, and analyzed via flow cytometry to evaluate cell death during HCoV-OC43 or mock infection. Mean with range is plotted for technical replicates.

